# p.E152K-STIM1 mutation deregulates Ca^2+^ signaling contributing to chronic pancreatitis

**DOI:** 10.1101/2020.01.22.916254

**Authors:** Miguel Burgos, Reginald Philippe, Fabrice Antigny, Paul Buscaglia, Emmanuelle Masson, Sreya Mukherjee, Pauline Dubar, Cédric Le Maréchal, Florence Campeotto, Nicolas Lebonvallet, Maud Frieden, Juan Llopis, Beatriz Domingo, Peter B. Stathopulos, Mitsuhiko Ikura, Wesley Brooks, Wayne Guida, Jian-Min Chen, Claude Ferec, Thierry Capiod, Olivier Mignen

## Abstract

Since deregulation of intracellular Ca^2+^ can lead to intracellular trypsin activation and STIM1 (stromal interaction molecule-1) protein is the main regulator of Ca^2+^ homeostasis in pancreatic acinar cells, we explored the Ca^2+^ signaling in 37 STIM1 variants found in three pancreatitis patient cohorts. Extensive functional analysis of one particular variant, p.E152K, identified in three patients, provided a plausible link between dysregulated Ca^2+^ signaling within pancreatic acinar cells and chronic pancreatitis susceptibility. Specifically, p.E152K, located within the STIM1 EF-hand and sterile α-motif domain, increased the release of Ca^2+^ from the endoplasmic reticulum in patient-derived fibroblasts and transfected HEK293T cells. This event was mediated by altered STIM1-sarco/endoplasmic reticulum calcium transport ATPase (SERCA) interactions and enhanced SERCA pump activity leading to increased Store Operated Calcium Entry (SOCE). In the pancreatic AR42J cells expressing the p.E152K variant, Ca^2+^-signaling perturbations correlated with defects in trypsin activation and secretion, and increased cytotoxicity after cholecystokinin stimulation.

**Summary statement:** p.E152K-STIM1 variant found in pancreatitis patients leads to intracellular changes in calcium homeostasis through SERCA interaction, enabling intracellular trypsin activation and pancreatic acinar cell death.

## INTRODUCTION

Cell damage observed in experimental acute pancreatitis results in part from abnormal intracellular calcium concentrations ([Ca^2+^]_i_) (Petersen, 2009; Gerasimenko et al., 2014). Elevated [Ca^2+^]_i_ signals can be elicited by combinations of fatty acids, alcohol or bile acids, leading to the appearance of fatty acid ethyl esters, which mediate the toxic alcohol effects on pancreatic acinar cells. This higher [Ca^2+^]_i_ results from the excessive release of endoplasmic reticulum (ER) Ca^2+^ stores or from an increase in Ca^2+^ influx such as store-operated calcium entry (SOCE) regulated by stromal interaction molecule-1 (STIM1) and mediated by ORAI1 proteins. The sustained Ca^2+^ signals and cytosolic Ca^2+^ overload contribute to the development of acute pancreatitis (Gerasimenko et al., 2009; Lur et al., 2011; Wen et al., 2015) through initiating activation of intracellular proteases responsible for cell autodigestion (Raraty et al., 2000; Gerasimenko et al., 2013; Zhu et al., 2018). Actually, the inhibition of calcium channel activity has been described as protective in pancreatitis and represents a therapeutic option (Son et al., 2019; Waldron et al., 2019).

STIM1 is a single-pass transmembrane protein mainly localized to the ER membrane and has been established as the main ER Ca^2+^ sensor in non-excitable and excitable cells (Yuan et al., 2009). The EF-hand and sterile α-motif (EF-SAM) domains of STIM1 are located in the luminal region of the ER and together act as a Ca^2+^ sensor to initiate SOCE activation. After ER Ca^2+^ store depletion, STIM1 oligomerizes and subsequently translocates to ER-plasma membrane junctions (Liou et al., 2005; Roos et al., 2005) allowing interaction with store-operated Ca^2+^ channels (Wu et al., 2006; Park et al., 2009; Nwokonko et al., 2017). The association between *STIM1* gene variants and the development of different diseases has been elucidated. Among them, we can find *STIM1* variants driving to loss of function in autoimmunity and immunodeficiencies like E136X (Picard et al., 2009) and homozygous p.L74P and p.L374P (Parry et al., 2016; Vaeth et al., 2017), or reporting a gain of function like p.D84G/p.H84N/p.H109R for tubular-aggregate myopathy (Böhm et al., 2013), p.R304W for Stormorken syndrome (Misceo et al., 2014; Morin et al., 2014; Nesin et al., 2014), p.I115F (Hedberg et al., 2014) and p.D84E (Noury et al., 2017) for tubular-aggregate myopathy, and p.S88G/p.R304Q for neuromuscular malfunctions (Harris et al., 2017). The importance of Ca^2+^ signaling for the regulation of pancreatic zymogen activation and the key role of STIM1 in this process suggests that variants in the *STIM1* gene may also contribute to chronic pancreatitis by disturbing Ca^2+^ homeostasis within the pancreatic tissue. In this regard, Sofia and colleagues have recently analyzed the *STIM1* in 80 patients with idiopathic chronic pancreatitis (ICP) and found three missense mutations in different patients (Sofia et al., 2016). Indeed, a recently genetic study of the Ca^2+^ channel TRPV6 in pancreatitis cohorts described variants of the protein linked to Ca^2+^ deregulation leading to the appearance of pancreatitis features (Masamune et al., 2020).

In this article, we analyzed the calcium signaling of 37 STIM1 variants found in 3 pancreatitis patient cohorts containing 2057 patients and 3322 controls (Masson et al 2019, Genetic analysis of the STIM1 gene in chronic pancreatitis, doi: http://dx.doi.org/10.1101/691899). Extensive functional analyses of a particular variant, E152K, suggested that Ca^2+^ deregulation within the pancreatic acinar cells may potentially modify the risk of chronic pancreatitis.

## RESULTS

### Identification of *STIM1* variants in ICP patients

As described, the 37 distinct variants found in a cohort of chronic pancreatitis patients were not evenly distributed along the STIM1 amino acid sequence, approximately 65% (*n* = 24) of them occurring in the last 185 residues (27%) of the 685-amino acid protein (Fig. 1A). Whereas the 24 variant C-terminal cluster is located primarily within the poorly characterized region of STIM1 between CC3 and P/S, a small cluster comprising four variants is located between the signal peptide (S) and the first EF-hand domain (cEF). By contrast, another small cluster comprising five variants is located within a well-defined region, the SAM domain (Fig 1A).

**Figure 1.**
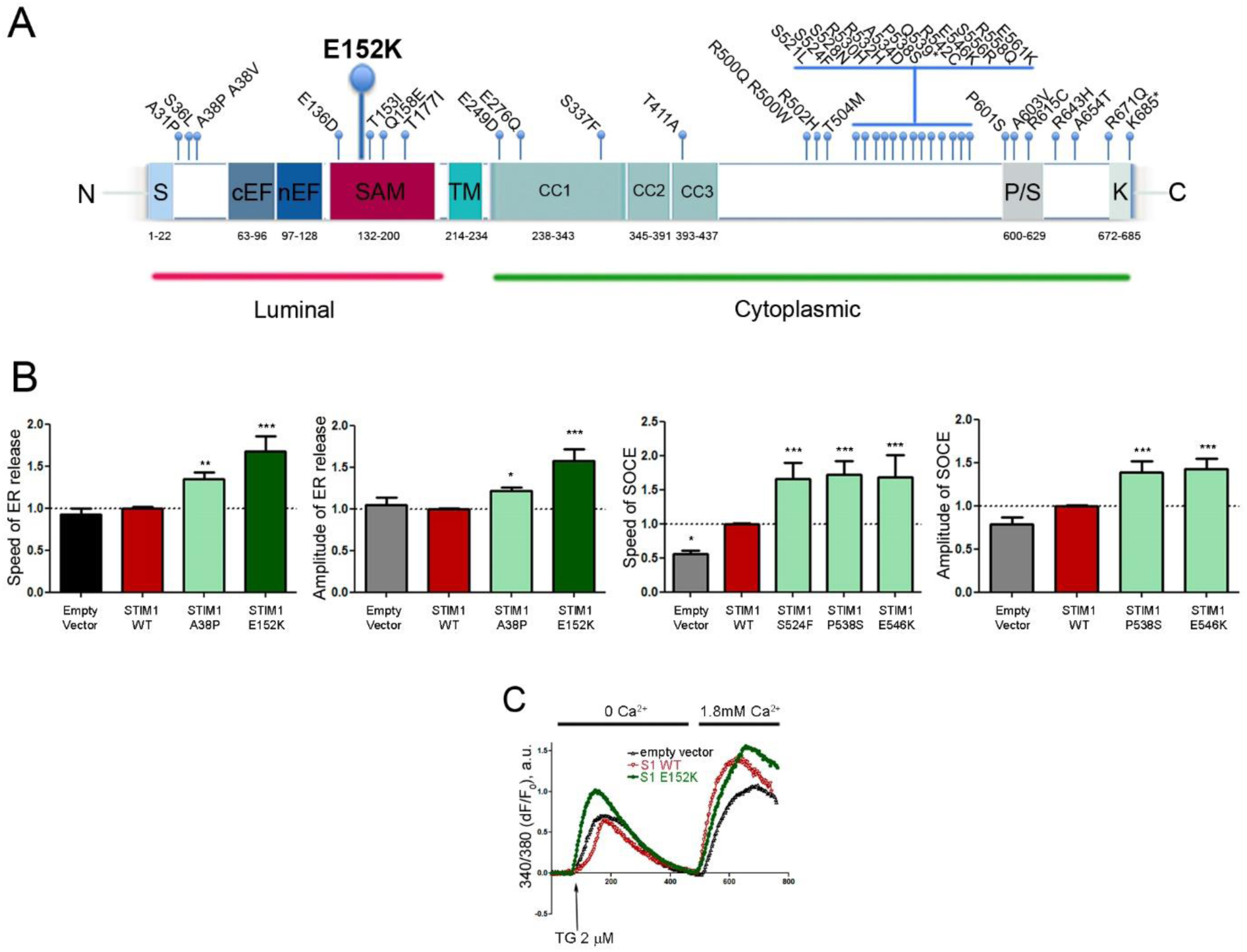
Functional characterization of *STIM1* variants in HEK293T cells. (**A**) Domain structure of the human STIM1 protein and spatial distribution of all 37 *STIM1* variants identified in this study. S, signal peptide; cEF and nEF hands, canonical and non-canonical EF hands; TM, transmembrane region; CC1, CC2 and CC3, coiled-coil domains 1, 2 and 3; P/S, proline/serine enriched domains; K, lysine-enriched domain. (**B**) Bar graphs showing the frequencies for the parameters speed and amplitude of ER Ca^2+^ release, and speed and amplitude of SOCE in HEK293T cells transfected with the specified *STIM1* variants and normalized to STIM1-WT. (**C**) Representative traces of TG-evoked Ca^2+^ transients in Fura-2-loaded HEK293T cells expressing WT STIM1, STIM1 E152K or an empty vector.

### Functional screening of all 37 *STIM1* variants in HEK293T cells

To explore the effects of *STIM1* variants on Ca^2+^ flux, *STIM1* variants were transiently expressed in HEK293T cells. The speed and amplitude of Ca^2+^ release from ER stores and SOCE after Thapsigargin (TG) addition were then evaluated 48 h after transfection in fura-2 loaded cells (Table S1). As expected, over-expression of wild-type (WT) STIM1 induced an increase in the SOCE rate as compared to cells expressing an empty vector without significantly altering ER Ca^2+^ release (Fig. 1B). Significant increases in SOCE were also observed in cells expressing p.S524F, p.P538S and p.E546K *STIM1* variants in comparison to WT STIM1 (Fig. 1B). It should however be noted that two of the three variants, p.S524F and p.E546K, were found in both patients and controls with equal or comparable frequencies (Masson et al 2019, Genetic analysis of the STIM1 gene in chronic pancreatitis, doi: http://dx.doi.org/10.1101/691899).

Surprisingly, ER Ca^2+^ release induced by SERCA pump inhibition by TG was significantly increased in cells expressing STIM1 p.A38P and p.E152K compared to WT STIM1-expressing cells (Fig. 1B,C). We observed the same effect after tBHQ addition with respect to STIM1 p.E152K by comparison with WT STIM1-expressing cells (Fig. S1A).

The p.E152K variant was of particular interest for three reasons. First, it was observed in three patients but not in controls (Table S1). It should also be pointed out that p.E152K is extremely rare in the general population; its average allele frequency in gnomAD (gnomad.broadinstitute.org/) was 0.0001162 (30/258080; as of November 30, 2018). Second, p.E152K affects a highly conserved amino acid of the EF-SAM domain located within the 7th α-helix (Fig. S1B). The highly conserved EF-SAM domain is central to STIM1 oligomerization (Stathopulos et al., 2006). Gain of function variants in the EF-hand domain have been previously reported to induce constitutive STIM1 oligomerization and have been associated with tubular-aggregate myopathy (Böhm et al., 2013; Hedberg et al., 2014; Lacruz and Feske, 2015). Third, and somewhat intriguingly, all three patients harboring the *STIM1* p.E152K variant were found to harbor heterozygous variants in other known chronic pancreatitis susceptibility genes. Thus, of the two French patients, one harbored *PRSS1* p.N29I whereas the other harbored *PRSS1* p.P36R. Both *PRSS1* p.N29I and *PRSS1* p.P36R are rare variants whereas *PRSS1* p.N29I is a well-known pathogenic mutation; *PRSS1* p.P36R is currently classified as benign (see the Genetic Risk Factors in Chronic Pancreatitis Database; http://www.pancreasgenetics.org/index.php). In the Chinese patient harboring the *STIM1* p.E152K variant, the situation is more complicated: the patient also harbored the *SPINK1* c.194+2T>C variant, the *PRSS1* c.623G>C (p.G208A) variant and the *CTRC* c.180C>T (p.G60=) variant; all three of the latter variants have been recently shown to be associated with chronic pancreatitis in a large Chinese cohort study (Zou et al., 2018).

In view of the aforementioned points, we focused our functional exploration on the p.E152K variant.

### Ca^2+^ signaling deregulation in fibroblasts obtained from an ICP patient harboring the p.E152K variant

Fibroblasts were obtained from the patient carrying the *STIM1* p.E152K variant and the *PRSS1* p.P36R variant (II.2), his mother who harbored neither of the two variants (I.2) and his brother (II.1) carrying only the *PRSS1* p.P36R variant (Fig. 2A). Clear and significant increases in the speed and amplitude of the ER Ca^2+^ release as well as the SOCE rate were observed in fibroblasts from the patient as compared to the two aforementioned relatives (Fig. 2B).

**Figure 2.**
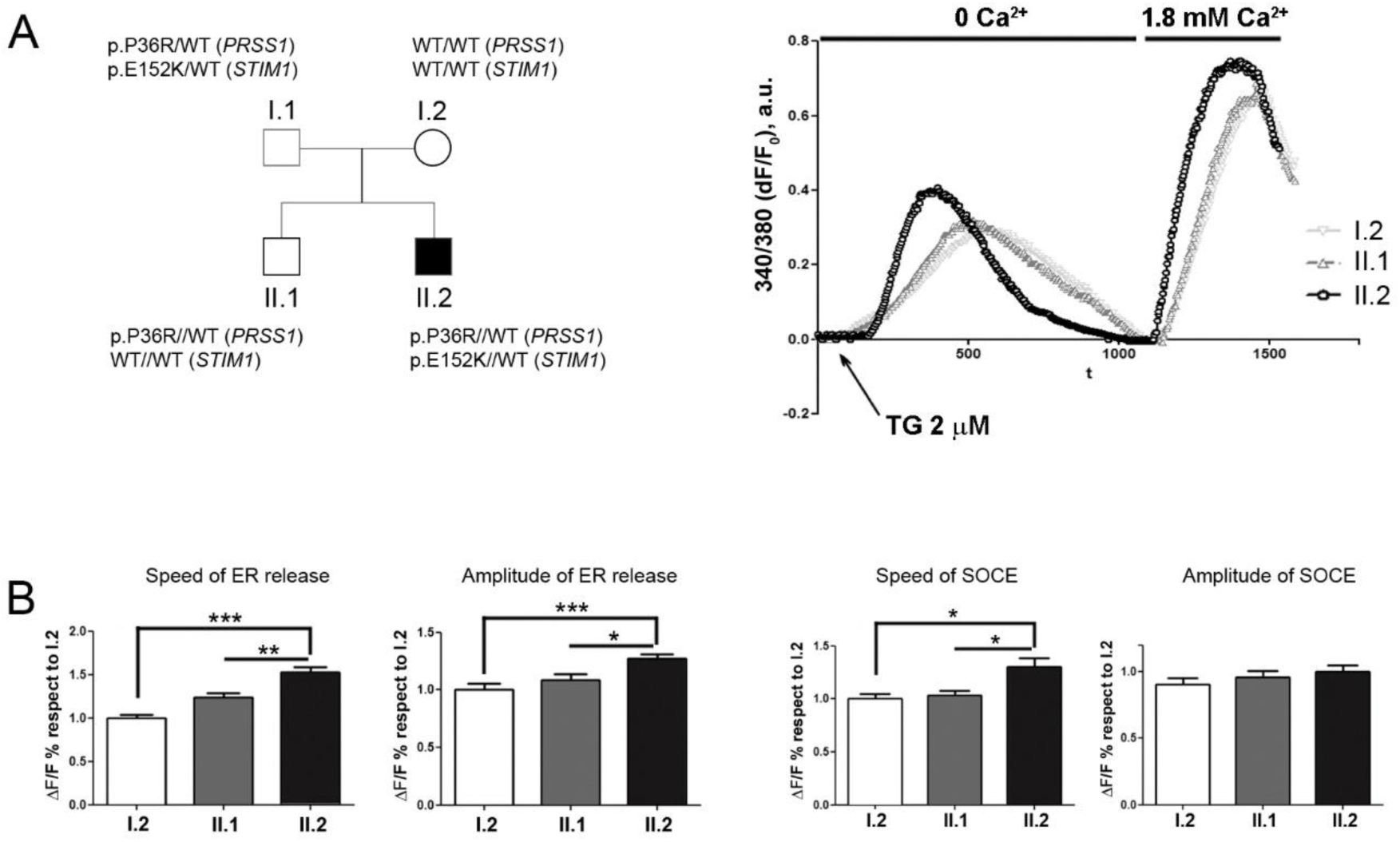
Analysis of Ca^2+^ responses to TG in fibroblasts isolated from a chronic pancreatitis patient harboring the *STIM1* p.E152K variant**. (A)** Left panel, family tree of the patient (i.e., II.2). *STIM1* genotypes are given for each family member. Right panel, representative traces of TG-evoked Ca^2+^ transients in Fura-2-loaded fibroblasts from patient II.2 and two of his relatives (I.1 and II.1). (**B)** Average values of the rate and amplitude of ER Ca^2+^ release and SOCE normalized to I.1 parameter values.

### Modulation of ER Ca^2+^ homeostasis in HEK293T cells expressing the p.E152K variant

One possible explanation for the increased ER Ca^2+^ release is enhanced passive Ca^2+^ leakage from the ER when TG blocks SERCA. However, receptor-mediated Ca^2+^ release consequent to the activation of InsP_3_ receptors (InsP_3_Rs) following carbachol (CCh) stimulation (Fig. 3A and Fig. S1C, left panel) or ATP (Fig. 3B and Fig. S1C, right panel) is also significantly enhanced. A series of p.E152K/D/V/Q/R variants introduced into STIM1 confirmed the key role of the negative charge contributed by this basic amino acid in relation to the enhanced ER Ca^2+^ release phenotype (Fig. S1D). For example, p.E152R shows the same ER Ca^2+^ increase as p.E152K.

**Figure 3.**
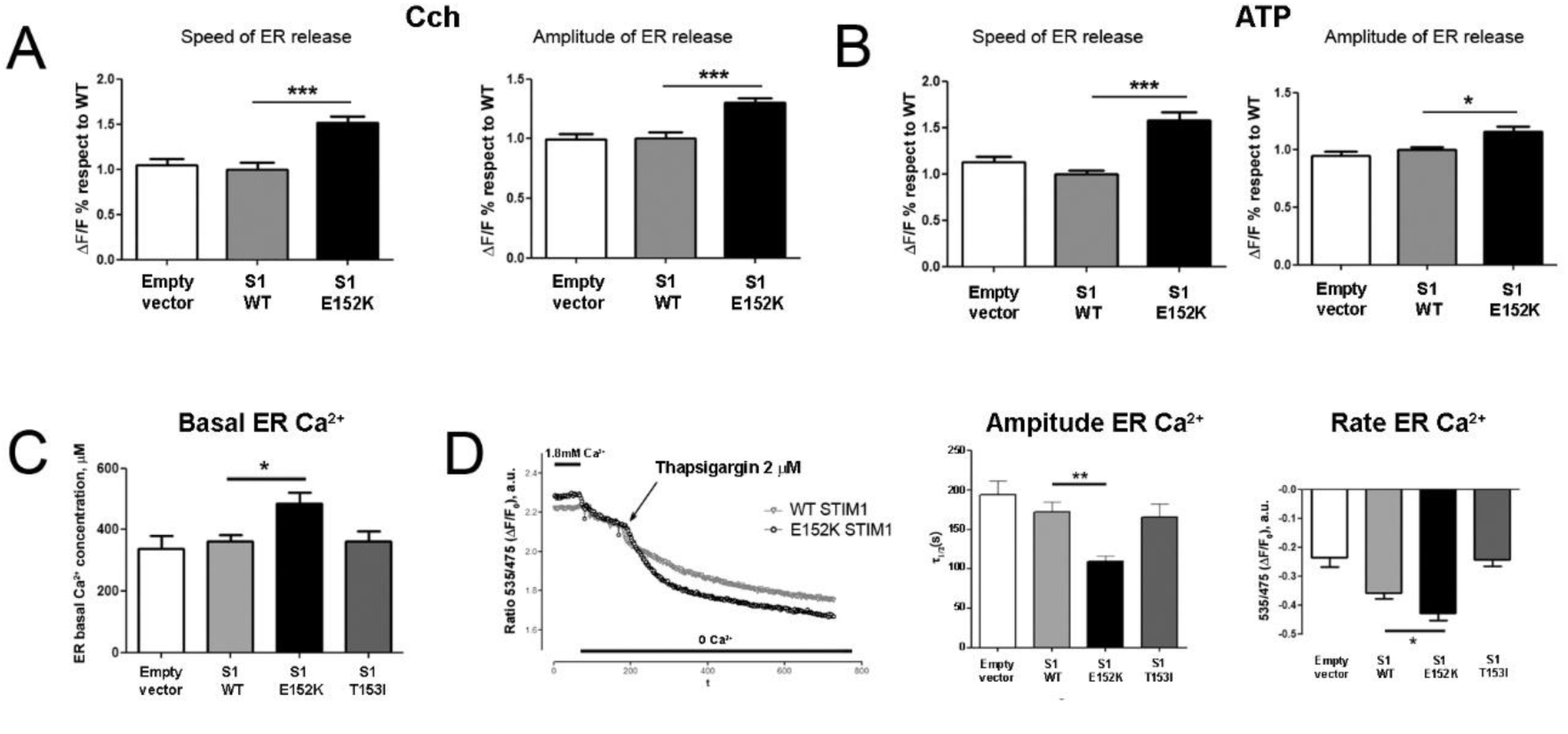
Analysis of the Ca^2+^ response in HEK293 cells expressing either WT or mutated STIM1. **(A)** Average values of the speed (left panel) and amplitude (right panel) of ER Ca^2+^ release after 100 µM CCh treatment in the absence of extracellular Ca^2+^. (**B)** Average values of the speed (left panel) and amplitude (right panel) of ER Ca^2+^ release after 100 µM ATP treatment in the absence of extracellular Ca^2+^. (**C**) Basal [Ca^2+^]_er_ (µM) evaluated using the D1ER probe and values normalized based on the calibration curve. (**D**) Representative traces of [Ca^2+^]_er_ variation after addition of 1 µm TG (Ca^2+^-free medium) to D1ER-transfected HEK293T cells expressing WT STIM1 or E152K STIM1 (left hand panel). Center panel, quantification of the ER Ca^2+^ release rate (_1/2_) in cells expressing WT STIM1, E152K STIM1, T153I STIM1 or empty vector. Right hand panel, quantification of the amplitude of [Ca^2+^]_ER_ release after TG (1µM) treatment in cells expressing WT STIM1, E152K STIM1, T153I STIM1 or empty vector, n = 42 (empty vector), 106 (WT), 76 (E152K) and 54 (T153I).

Next, we directly monitored changes in ER Ca^2+^ concentration in HEK293T cells transfected with the ER-targeted cameleon probe D1ER (Palmer et al., 2004). ER Ca^2+^ concentration ([Ca^2+^]_er_) was determined after D1ER titration (Fig. S1E) as previously described (Shen et al., 2011). The simplest explanation for the increase in ER Ca^2+^ release is that a higher [Ca^2+^]_er_ enhances the driving force for the ER Ca^2+^ efflux. No significant increase in [Ca^2+^]_er_ was observed when over-expressing WT STIM1 or STIM1-T153I (Fig. 3C). However, ER Ca^2+^ concentration was markedly increased in cells expressing STIM1-E152K as compared to WT STIM1 (Fig. 3C, 484 vs 363 μM, *P* < 0.05). After blocking SERCA using TG, a significant enhancement (*P* < 0.001) of both the rate (36%) and amplitude (23%) of [Ca^2+^]_er_ decrease was only observed in STIM1-E152K-expressing cells (Fig. 3D).

### Interaction between SERCA and STIM1 modulates SERCA activity and ER refilling

An increase in SERCA pump activity would explain the increase in [Ca^2+^]_er_ and Ca^2+^ efflux in cells expressing STIM1-E152K. SERCA pump activity, estimated by quantifying ER Ca^2+^ refilling following store depletion with tBHQ in permeabilized and non-permeabilized D1ER-expressing cells (Fig. 4A,B), is clearly enhanced in STIM1-E152K-expressing cells independently of SOCE as demonstrated by experiments performed under permeabilizing conditions.

**Figure 4.**
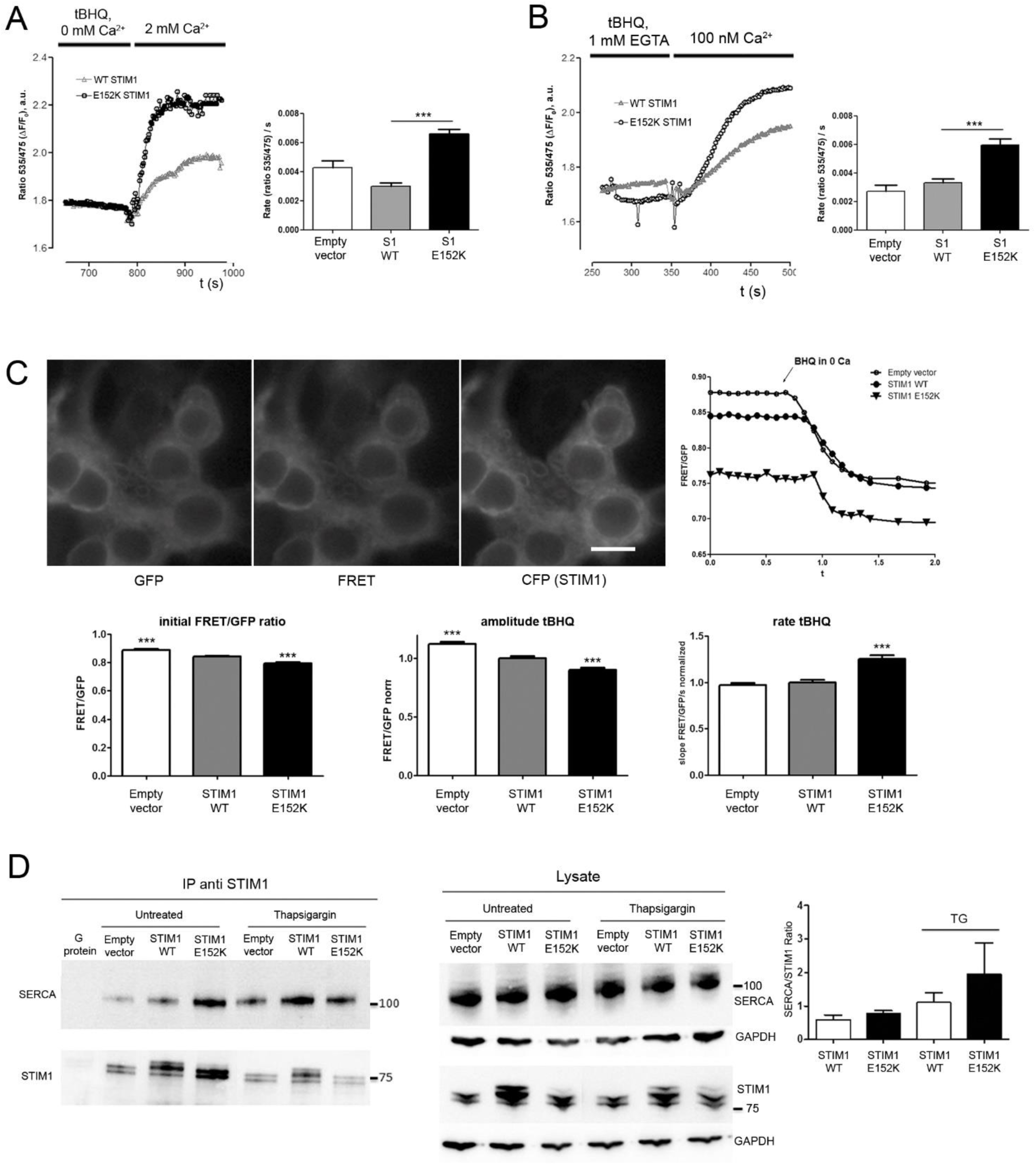
STIM1-SERCA modification induced by the E152K variant. **(A, B)** Cells were transfected with either WT or E152K STIM1; ER Ca^2+^ changes were evaluated in cells transfected with ER-targeted cameleon probe D1ER. In (**A**), ER Ca^2+^ depletion was induced by 15 µM tBHQ, and ER Ca^2+^ refilling was evaluated by changing the Ca^2+^-free extracellular medium using a solution containing 1.8 mM Ca^2+^ (representative traces in left hand panel) and quantification of the ER Ca^2+^ refilling rate in cells expressing WT STIM1, E152K STIM1 or an empty vector (right hand panel). n = 19 (empty vector), 67 (WT) or 46 (E152K) (right hand panel). In (**B**), the quantification of ER Ca^2+^ refilling rate in cells expressing WT STIM1, E152K STIM1 or an empty vector was performed under permeabilizing conditions. Left hand panel: representative traces of ER Ca^2+^ variations in cells permeabilized by 60 µM digitonin and after addition of 15 µM tBHQ. ER Ca^2+^ refilling was induced by addition of 100 nM Ca^2+^. n = 8 (empty vector), 91 (WT) or 71 (E152K) (right hand panel). (**C**) Images showing a double transfection of HEK293 cells using a FRET-based intramolecular SERCA construct (GFP-tagRFP) and STIM1 WT-CFP (or alternatively E152K-STIM1-CFP). Scale bar, 50 µM. Left panel, fluorescence images for GFP, FRET and STIM1-CFP. A basal FRET/GFP fluorescence ratio level is observed, which decreases as the SERCA adopts an open conformation after tBHQ addition (right panel). Histograms show the quantification of differences in basal FRET/GFP ratio (left subpanel), amplitude of tBHQ response (center) and rate of tBHQ response (right subpanel). (**D**), HEK293 cells transfected with WT STIM1, E152K STIM1 or an empty vector were left untreated or treated with 2 µM TG in 1.8 mM Ca^2+^-solution for 5 min and were subjected to immunoprecipitation with anti-STIM1 antibody (lanes 1 to 6). Eluted fractions after immunoprecipitation (IP) (left hand panel) and lysates (right hand panel) were analyzed by SDS-PAGE and Western blot with anti-SERCA- and anti-GOK/STIM1-specific antibodies. Right panel, quantitative analysis of STIM1/SERCA interaction from 7 independent experiments. Data are presented as average intensity values of SERCA bands normalized to the respective STIM1 band intensity in a specific IP.

Hence, we hypothesized that SERCA activity can be directly regulated by STIM1. Different studies have reported the interaction of STIM1 with SERCA2 and SERCA3 pumps (López et al., 2008; Sampieri et al., 2009; Manjarrés et al., 2010). Using a FRET-based SERCA construct (Satoh et al., 2011; Hou et al., 2012; Pallikkuth et al., 2013), we sought to establish whether the stronger association between SERCA and STIM1-E152K could modulate SERCA conformational state changes. At rest, a basal FRET level is seen, whereas after tBHQ addition, the FRET signal decreases as the SERCA cytoplasmic headpiece is driven to an “opened” conformation (Hou et al., 2012). Remarkably, we observed a reduction in the basal FRET/GFP ratio in cells over-expressing WT STIM1 in comparison to empty vector, and an even lower ratio in cells expressing E152K-STIM1 (Fig. 4C). After tBHQ addition, the amplitude and rate of SERCA conformational change is higher in E152K expressing cells than WT. The maximal rate of FRET/GFP decrease induced by Ca^2+^ store release was significantly enhanced in cells expressing STIM1-E152K. In terms of amplitude, both WT STIM1 and STIM1-E152K diminished the FRET/GFP decrease induced by tBHQ, E152K having a stronger effect. When ER Ca^2+^ was depleted, SERCA displayed a more rapid conformational change in the presence of STIM1-E152K in comparison to WT STIM1, although smaller in amplitude.

To reinforce our hypothesis of a modification of STIM1/SERCA interaction in the presence of E152K variant, we performed immunoprecipitation experiments. STIM1 and SERCA co-immunoprecipitate under resting conditions and the association is increased after store depletion when WT STIM1 is overexpressed (Fig. 4D). Interestingly, an increase in SERCA/STIM1 co-immunoprecipitation was observed in STIM1-E152K-expressing cells as compared to cells expressing WT STIM1 after TG treatment (Fig. 4D). Taken together, these results suggest that under resting [Ca^2+^]_er_ conditions, a fraction of STIM1 is already associated with SERCA. When [Ca^2+^]_er_ is lowered, an enhanced functional association between the STIM1-E152K and SERCA compared to WT STIM1 may then increase this Ca^2+^ pump activity and modulate its Ca^2+^ affinity resulting in a higher [Ca^2+^]_er_.

Taken together, these data suggest that the p.E152K variant modifies the degree of association between SERCA and STIM1. Among other explanations, the greater STIM1-E152K association with SERCA before store depletion could favour an intermediate cytoplasmic conformational state which responds faster after Ca^2+^ depletion.

### Intrinsic biophysical properties of STIM1-E152K

It is now well established that the EF-SAM domain plays a critical role in the regulation of STIM1 oligomerization (Stathopulos et al., 2006). Computational modelling of STIM1 for both the WT and the p.E152K variant to discern possible changes in the mutated STIM1 structure (Supplementary note and Fig. S2) suggest that a difference in local charge due to the p.E152K mutation may affect STIM1 protein-protein interactions. Using a FRET approach, we observed a difference in FRET multimerisation between WT and E152K mutated STIM1. We observed a significantly higher FRET level between CFP-STIM1-E152K and YFP-STIM1-E152K by comparison with the CFP-WT STIM1/YFP-WT STIM1 constructs, consistent with an increased efficiency of STIM1-E152K homo-oligomerization (Figure 5a). FRET analysis of WT-STIM1/E152K mutated STIM1 multimerization as expected in an heterologous expression condition, is not relevant due to the higher homo-oligomerization efficiency of mutated STIM1 compared to WT STIM1 that will bias CFP-WT/YFP-E152K FRET data interpretation.

**Figure 5.**
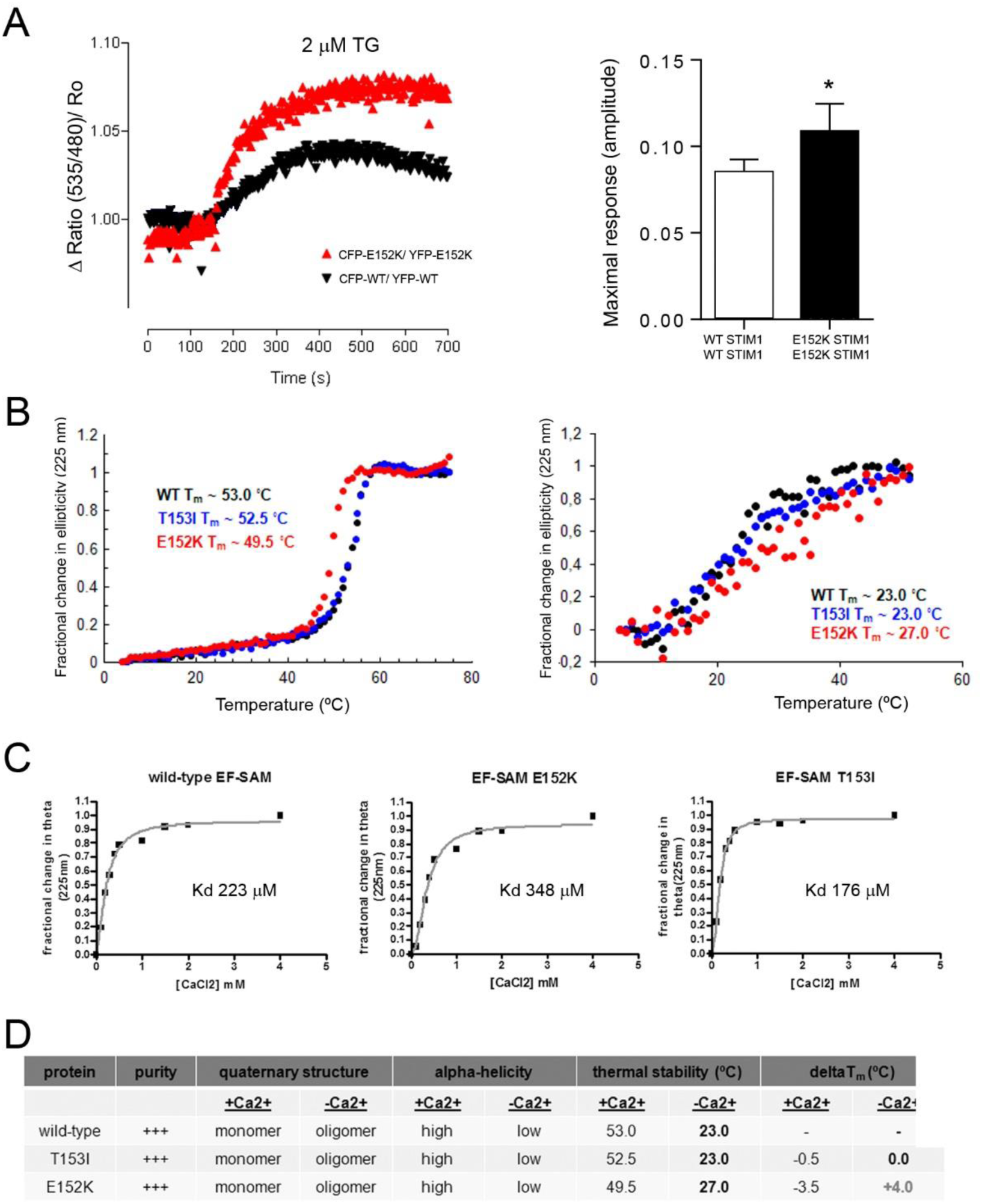
Experiments on the biophysical properties of STIM1-E152K. (**A**) Left hand panel; representative traces of changes in FRET signal (CFP-STIM1 WT / YFP-STIM1 WT, and CFP-STIM1 E152K/YFP-STIM1 E152K) were measured in transfected HEK cells after addition of 1 µM TG to induce ER Ca^2+^ store depletion. (**B**) Thermal stability of WT and mutant EF-SAM domains. Thermal stabilities of WT EF-SAM (black circles), T153I EF-SAM (blue circles) and E152K EF-SAM (red circles) domains in the presence (left) and absence (right) of 5mM CaCl_2_ were estimated from the changes in far-UV-CD at 225nm as a function of temperature. The apparent midpoint of temperature denaturation (Tm) was defined as the temperature where the fractional change in ellipticity was 0.5. Protein concentrations were 10 µM and 40 µM in the Ca^2+^-loaded and – depleted states, respectively. (**C**) Ca^2+^ binding affinity estimates of WT and mutant EF-SAM domains. Binding curves of 40 µM protein were acquired as the fractional change in ellipticity as a function of CaCl_2_ concentration at 20°C. Data were fitted to the Hill equation, assuming a negligible effect of protein concentration on the dissociation constant (Kd) since [protein]<<Kd. Buffers in (B) and (C) were 20mM Tris, 150mM NaCl, pH 8. (**D**) Summary of WT and mutant EF-SAM biophysical characteristics.

Biophysical analyses were then performed on purified EF-SAM domains of WT or mutated human STIM1 (Fig. S3A). Far-UV circular dichroism (CD) experiments confirmed that the mutant EF-SAM domains retain the ability to sense Ca^2+^ changes through a marked alteration in structure, consistent with the ability of full-length STIM1 E152K to signal SOCE after ER Ca^2+^ store depletion (Fig. S3B). Thermal stability studies of the EF-SAM domains revealed that the E152K variant destabilized the Ca^2+^-bound EF-SAM structure (T_m_ WT = 53°C versus T_m_ E152K = 49.5°C, Fig. 5B *left subpanel*). In contrast, the E152K variant appeared to stabilize the Ca^2+^-depleted EF-SAM structure (Tm WT ∼ 23°C versus Tm E152K ∼ 27°C, Figure 5b right subpanel), with the caveat that both the WT and E152K data showed a high variability due to the overall instability. These results suggest that the enhanced multimerization displayed by E152K STIM1 as compared to WT may be due to the destabilized Ca^2+^-loaded state which more readily undergoes a transition to the oligomerization-prone Ca^2+^-depleted state and/or a modified Ca^2+^-depleted state with enhanced oligomer stability.

Size exclusion chromatography with in-line multiangle light scattering (SEC-MALS) allowed us to conclude that in the presence of Ca^2+^, E152K EF-SAM exists primarily as a monomer, but undergoes a shift in the self-association equilibrium towards dimers and oligomers in the absence of Ca^2+^ as observed for WT STIM1 and T153I EF-SAM (Fig. S3C). No significant change in the Ca^2+^ binding affinity of E152K STIM1 was observed and the measured affinities were in the same range as those previously reported for WT STIM1 EF-SAM (Stathopulos et al., 2006) (Fig. 5D). Taken together, these data suggest that the E152K EF-SAM variant displays WT-like Ca^2+^ sensitivity characteristics in terms of Ca^2+^ binding, secondary structure and oligomerization. However, changes in the stability of STIM1 E152K EF-SAM with and without Ca^2+^ may contribute to aberrant ER Ca^2+^ homeostasis and differences in STIM1 protein interactions. The biophysical properties of the different EF-SAM domains are summarized in Fig. 5D.

### Functional consequences of E152K STIM1 in rat pancreatic acinar cells

In order to probe the physiological consequences of STIM1-E152K expression with a view to potentially linking the observed defects in Ca^2+^ signaling to pancreatitis, we next evaluated, in rat pancreatic AR42J cells, the effects of the STIM1 p.E152K variant on Ca^2+^ signaling, trypsin activation, trypsinogen secretion and the subsequent consequences for cell fate. As observed in HEK293T cells, a significant increase in the rate and amplitude of ER Ca^2+^ release induced by agonist stimulation with cholecystokinin (CCK) (Fig. 6A) or Cch (Fig. S4A) or by TG (Fig. S4B) was obtained in AR42J cells expressing STIM1-E152K (Fig. S4C).

**Figure 6.**
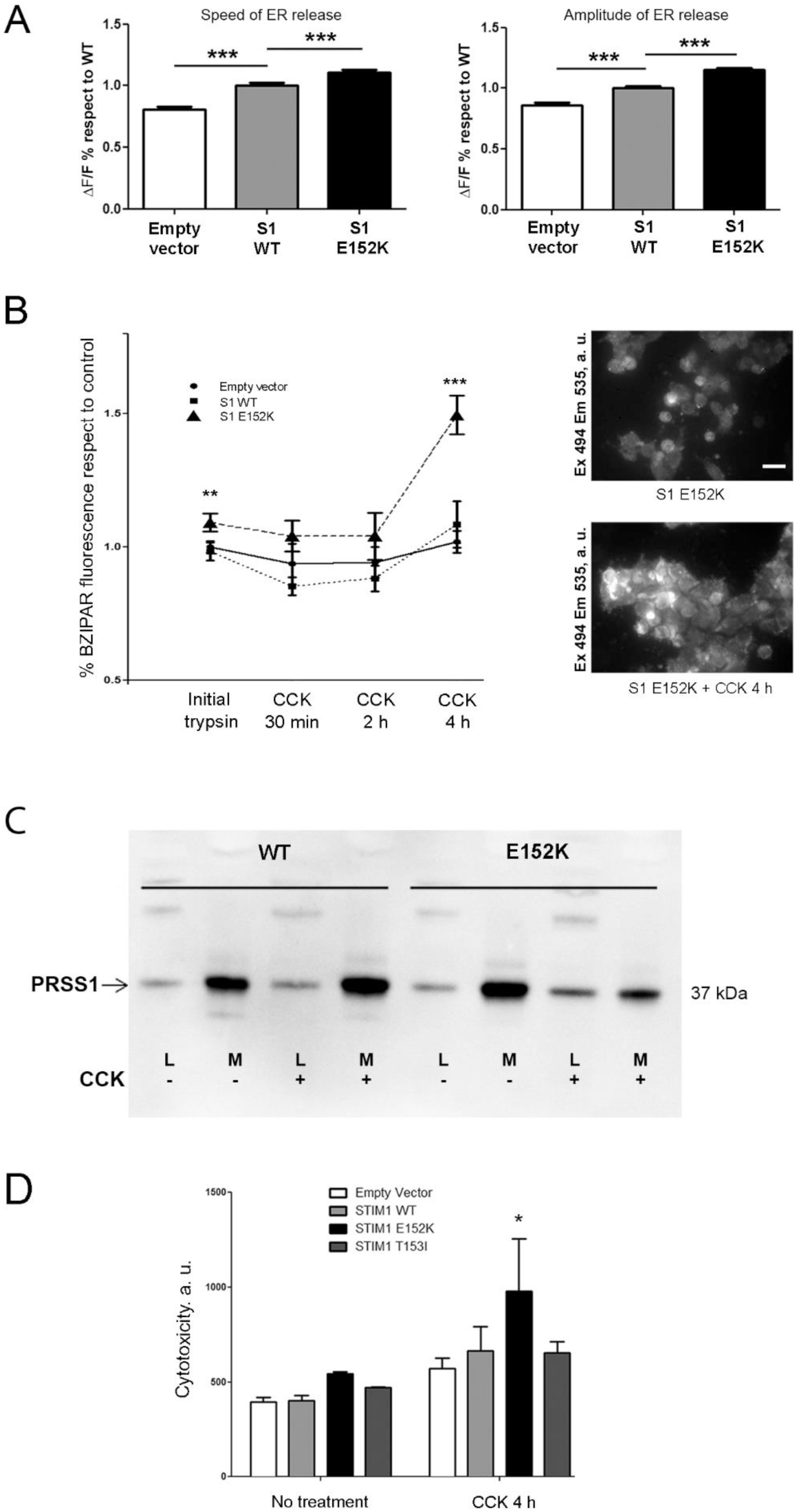
Functional consequences of E152K STIM1 in rat pancreatic acinar cells. **(A)** Average values of the speed (left hand panel) and amplitude (right hand panel) of ER Ca^2+^ release induced by the addition of 1 nM CCK in AR42J cells transfected with E152K STIM1, WT STIM1 or an empty vector. Values were normalized to WT STIM1. n = 449 (empty vector), 705 (WT) or 512 (E152K). (**B**) Left hand panel, quantification of BZiPAR fluorescence at different time points in cells expressing WT STIM1, E152K STIM1 or empty vector after CCK treatment (1 nM). Right hand panel, representative images of BZiPAR fluorescence in E152K STIM1-expressing cells, before and after 4h of CCK treatment. Cells were excited at 494 nm for 150 ms, and fluorescence emission was collected at 535 nm. Scale bar, 50 µM. (**C**) Analysis by Western blot of PRSS1 intracellular expression and its presence in the conditioned medium before and after stimulation with CCK (1 nM) of cells expressing E152K or WT STIM1. (**D**) Quantification over time of fluorescence signal evaluating cytotoxicity (celltox) in cells transfected with E152K STIM1, T153I STIM1, WT STIM1 or an empty vector after CCK addition (1 nM). Cell toxicity was measured using the CellTox™ Green Cytotoxicity Assay kit (Promega), and results were normalized to living cell numbers as evaluated with the CellTiter 96^®^ AQueous One Solution Cell Proliferation Assay Kit (Promega) following the manufacturer’s instructions.

Intra-acinar activation of zymogens is a key event in the initiation of acute pancreatitis (Krüger et al., 2000) and the increased Ca^2+^ mobilization observed in STIM1-E152K-expressing cells may lead to an increase in intracellular trypsin activation after stimulation. Trypsin activation was monitored before and after CCK stimulation using the BZiPAR probe (Raraty et al., 2000), which allowed us to detect constitutive activation of trypsin in AR42J cells that was enhanced following CCK stimulation in a time-dependent manner (Fig. 6B) but was prevented by pretreatment with benzamidine, a competitive inhibitor of trypsin (Fig. S4D). A higher amount of basal activated trypsin as well as a significant enhancement of trypsin activation after 4 h stimulation was detected in AR42J cells expressing STIM1-E152K as compared to WT STIM1 or empty vector (Fig. 6B) (12% increase in basal trypsin activity and 49% increase after 4h CCK treatment respectively, *P* < 0.001).

Another explanation proposed for cell degradation observed in pancreatitis is a decrease in Ca^2+^-dependent trypsinogen secretion (Gerasimenko et al., 2014). Trypsin cellular content and exocytosis were evaluated as previously described by Western blot of cell lysates and in conditioned media after CCK treatment of AR42J cells expressing PRSS1 (Kereszturi and Sahin-Tóth, 2009). An enhancement of intracellular trypsin and a decrease in trypsin secretion was observed in cells expressing STIM1-E152K as compared to cells transfected with WT STIM1 in this experiment after CCK treatment, suggesting a difference in trypsin secretion when *E152K* variant was present (Fig. 6C). In agreement with these findings, the basal level of toxicity was higher in cells expressing STIM1-E152K as compared to cells transfected with an empty vector, WT STIM1 or STIM1-T153I (Fig. 6D). An enhancement of cell toxicity following CCK stimulation was observed in AR42J cells expressing STIM1-E152K in comparison to WT STIM1, in agreement with the increase in trypsin auto-activation observed under the same conditions (Fig. 6B).

## DISCUSSION

Notably, none of the identified 37 rare *STIM1* variants identified in the 3 pancreatitis patients cohorts correspond to those previously reported to cause or predispose to other diseases (Lacruz and Feske, 2015), potentially strengthening the notion of the tissue-specific effects of different *STIM1* variants. However, functional screening of these variants in relation to the speed and amplitude of Ca^2+^ release from ER stores and SOCE failed to reveal any enrichment of functional variants in patients. This may reflect a common challenge that we face in the functional annotation of disease-predisposing or modifying variants that may either only subtly affect gene expression or protein structure and function, or alternatively may exert their effects in tissue- or cell type-dependent fashion. Nonetheless, extensive functional analysis of one particular variant, p.E152K, which was found in three patients (two French and one Chinese) but not in any of the 3,322 controls, established a plausible link between dysregulated Ca^2+^ signaling within the pancreatic acinar cells and chronic pancreatitis susceptibility. Since all three patients harboring the *STIM1* p.E152K variant also harbored variants in known chronic pancreatitis susceptibility genes in the trypsin-dependent pathway, the *STIM1* p.E152K variant may be acting as a disease modifier.

E152K is located in the EF-SAM domain, a region previously identified as being crucial for STIM1 function. So far, the role of the STIM1 EF-SAM domain has been uniquely associated with SOCE activation (Stathopulos et al., 2006; Stathopulos et al., 2008) whereas gain of function mutations in the EF-hand domain have only been associated with the constitutive activation of STIM1 and SOCE (Böhm et al., 2013; Lacruz and Feske, 2015; Noury et al., 2017). Here, we reveal a gain of function for the STIM1-E152K variant potentially caused by altered interactions with SERCA rather than enhanced EF-SAM oligomerization. This modification leads to deregulated calcium homeostasis through its interaction with SERCA and trypsin autoactivation, thereby triggering cell death. SERCA has been previously shown to interact with STIM1 and has been determined to be a fundamental component of the store-operated Ca^2+^ influx complex (SOCIC) formed by channels and proteins regulating SOCE and store refilling (Jousset et al., 2007; López et al., 2008; Manjarrés et al., 2010; Vaca, 2010).

The enhancement of SERCA pump activity after store depletion explains the increase in [Ca^2+^]_ER_ and Ca^2+^ release observed in STIM1-E152K-expressing cells. As proposed by Lopez and colleagues (López et al., 2008), STIM1 is already associated directly or indirectly with SERCA before store depletion under our experimental conditions. However, our study is the first to propose regulation of the SERCA pump by STIM1-mediated STIM1/SERCA physical interaction. We propose that the increased association between SERCA and STIM1-E152K, under resting conditions or after ER Ca^2+^ depletion, is responsible for the enhancement of SERCA pump activity and subsequently for the amplification of [Ca^2+^]_ER_ and ER Ca^2+^ release. An E152K variant favors the cytoplasmic “opened” state which occurs prior to the conformational change after Ca^2+^ depletion. Regulation of Ca^2+^ pump function via a physical interaction was previously demonstrated for the plasma membrane Ca^2+^ ATPase (PMCA) (Ritchie et al., 2012).

It is now clearly established that the cell damage observed in pancreatitis originates from an abnormal increase or maintained level of [Ca^2+^]_i_ (Petersen, 2009). Initial release of Ca^2+^ from both the ER and acidic pools in the apical region of pancreatic acinar cells is the critical starting point for the Ca^2+^-dependent activation of pro-enzymes such as trypsinogen. Expressing the STIM1-E152K variant in AR42J pancreatic cells clearly induces excessive Ca^2+^ release from the ER after cell stimulation that could be correlated with the cytotoxicity observed under these conditions, resulting from an increased basal trypsin activity along with decreased secretion of its trypsinogen precursor after cell stimulation. Regulation of cationic trypsinogen degradation and auto-activation along with trypsin inactivation are key protective mechanisms from pancreatic injury reported as being sensitive to Ca^2+^ (Szmola and Sahin-Tóth, 2007; Szabó et al., 2014).

In pancreatic acinar cells, the role of Ca^2+^ in zymogen granule secretion is well established (Messenger et al., 2014). The main role of ER Ca^2+^ release in the apical granular region of pancreatic acinar cells is to generate Ca^2+^ transients in local micro-domains to activate acinar fluid and enzyme secretion by exocytosis (Maruyama et al., 1993; Maruyama and Petersen, 1994; Petersen, 2015). Tightly regulated apical ER SERCA pump activity is therefore crucial for providing an optimal environment for controlled local Ca^2+^ signaling and regulated zymogen exocytosis. Excessive release of Ca^2+^ from the ER and subsequent perturbation in apical Ca^2+^ micro-domains have been linked to the major apical region structural and functional disorganization observed in pancreatitis such as the appearance of endocytic vacuoles in which trypsinogen has been transformed to active trypsin and mitochondrial depolarization (Gerasimenko et al., 2009; Petersen, 2015). Taken together, we surmise that the expression of some gain of function *STIM1* variants that enhance ER Ca^2+^ release may promote acinar cell damage thereby contributing to the etiology of chronic pancreatitis.

In summary, we have provided a number of pieces of evidence to support the view that a *STIM1* variant, p.E152K, could act as a disease modifier by causing specific changes in Ca^2+^ homeostasis that in turn increase intracellular trypsin activation and cell death. This finding may prompt new lines of research into ICP as well as opening new avenues for future therapeutic studies into the development of drugs designed to modulate Ca^2+^ signaling, with a view to preventing or at least delaying the transition from acute pancreatitis to chronic pancreatitis (Wen et al., 2015). Moreover, the identification of the *STIM1* p.E152K variant together with three known disease-predisposing variants (in three different trypsin-dependent pathway genes) in a same patient provided an exceptionally illustrative example for the complex etiology of chronic pancreatitis. Finally, this study led to the identification of a previously undescribed interaction between the STIM1 EF-SAM domain and SERCA, thereby generating new insights into the structure and function of STIM1.

## Material and Methods

### Reagents

Thapsigargin, cholecystokinin, carbachol, ATP, benzamidine, tBHQ, CCCP and digitonin were obtained from Sigma. Fura-2 BZiPAR and Protein G magnetic beads were from Invitrogen - Thermo Fisher Scientific (Waltham, MA). LipoD293 was from Tebu-bio (Le Perray-en-Yvelines, France). D1ER was kindly provided by N. Demaurex (Geneva, Switzerland). PRSS1 antibody was obtained from R&D systems (Minneapolis, MN), STIM1 antibody from BD Transduction Laboratories (Franklin Lakes, NJ), GAPDH antibody from Tebu-Bio, SERCA antibody from Santa Cruz Biotechnology (Dallas, TX) and anti-sheep secondary antibody was from R&D Systems. The ER-targeted cameleon probe D1ER was kindly provided by Drs. A. Palmer (La Jolla, CA) and R. Tsien (La Jolla, CA). YFP STIM1 and STIM1 CFP were generous gifts from Drs A. B. Parekh (Oxford, UK) and A. Rao (La Jolla, CA), respectively.

### Construction of STIM1 cDNA expression vector and mutagenesis

RNA was extracted with Trizol (Invitrogen, Carlsbad, CA) from Colo-357 cells. The SuperScript^®^ II Reverse Transcriptase, random hexamers (Qiagen, Courtaboeuf, France) and 2 µg RNA were used to synthesize first strand cDNA. RT-PCR was performed using the Phusion^®^ High-Fidelity DNA Polymerase (NEB, Evry, France) according to the manufacturer’s protocol in a 50-μl reaction mixture using forward primer 5’-ATGGATGTATGCGTCCGTCT-3’ and reverse primer 5’-CTACTTCTTAAGAGGCTTCT-3’. The PCR program comprised an initial denaturation at 98°C for 30 secs, followed by 40 cycles denaturation at 98°C for 20 secs, annealing at 62°C for 20 secs and extension at 72°C for 90 secs, with a final extension at 72°C for 5 min. After migration in a 1% agarose gel, the expected band (2,058bp) was purified using the MinElute PCR Purification Kit (Qiagen, Courtaboeuf, France). After adding the 3’ A-overhangs to the PCR purified product, the STIM1 cDNA was cloned into the pcDNA3.1/V5-His TOPO TA vector (Invitrogen, Cergy-Pontoise, France). All STIM1 variants were generated from the wild-type (WT) construct by site-directed mutagenesis using the QuikChange II XL Site-Directed Mutagenesis Kit (Stratagene, Massy, France). All resulting plasmids were checked by Sanger sequencing.

### Extraction of fibroblasts

Skin biopsies were washed in PBS, cut into small pieces and placed on 60 mm coated plates (Corning^®^ CellBIND^®^ Surface, Corning). Fibroblast expansion was made in DMEM completed with 10% FBS and 100 U/ml penicillin/streptomycin in a humidified incubator with 5% CO2 at 37°C. After the fibroblasts had colonized the plate, explants were removed, and fibroblasts expanded. At confluence, cells were trypsinized. Cells were used from passages 3 to 6, and placed on coverslips for image analysis.

### Cytosolic Ca^2+^ measurements

Cells were plated on 18 mm glass cover slips. Changes in cytosolic Ca^2+^ concentration were measured with Fura-2 (ThermoFisher, Waltham, MA). Cells were loaded with 4 µM Fura-2/AM for 45 min in the dark at room temperature in a medium containing 135 mM NaCl, 5 mM KCl, 1 mM MgCl2, 1.8 mM CaCl2, 10 mM Hepes, 10 mM glucose, pH 7.4. Cells were washed and equilibrated for 15 min in the same buffer to allow de-esterification of the dye. Ratiometric images of Ca^2+^ signals were obtained at room temperature using a microscope (IX71, Olympus) equipped with a monochromator illumination system (Polychrome V, TILL Photonics). Cells were illuminated alternately at 340/380 nm and emission was collected every 3 seconds through a 415DCLP dichroic mirror at 435 nm by a 14-bit CCD camera (EXiBlue, Qimaging). Image acquisition and analysis were performed with the Metafluor 6.3 software (Universal Imaging, West Chester, PA). The Excitation/Emission ratio was calculated for each cell. For normalization, the formula (F-F0)/F0, was used, where F is the Excitation/Emission ratio of each value and F0 is the initial ratio. The Ca^2+^-free solution contained 1mM EGTA instead of 1.8 mM CaCl2.

### ER Ca^2+^ measurements

HEK293T cells were transiently transfected using LipoD293 (tebu-bio) with 2 μg cDNA encoding the D1-ER construct 48 h before the experiments. Ratiometric images of Ca^2+^ signals were obtained using a microscope (Axio Observer, Zeiss) equipped with a Lambda DG4 illumination system (Sutter Instrument Company, Novato, CA) as previously described (Philippe et al., 2015). In situ calibration of the D1-ER was performed in medium containing 10 mM NaCl, 135 mM KCl, 1 mM MgCl2, 20 mM sucrose, 20 mM Hepes, 0.01 mM digitonin, 0.01 mM ionomycin, 0.005 mM CCCP (pH 7.1 with NaOH). 5 mM EGTA and 5 mM EDTA were used to buffer free [Ca^2+^] below 100 µM, and 10 mM EDTA to buffer free [Ca^2+^] between 100 µM and 1 mM (Max Chelator Winmaxc version 2.51). To evaluate the resting ER Ca^2+^ concentration, minimum and a maximum D1ER fluorescence values were measured at the end of each experiment. To quantify ER Ca^2+^ refilling, the slope of the increase of FRET signal was determined by a linear fit. To measure Ca^2+^ leak rates of the ER, passive ER depletion was induced by thapsigargin and the D1ER responses were fitted with a one-phase exponential decay function to extract the half-time (τ1/2).

Permeabilized HEK cells: after ER Ca^2+^ depletion by 15 µM tBHQ (15 min in Ca^2+^-free medium), cells were permeabilized by the addition of 60 µM digitonin (1 min). ER Ca^2+^ refilling was then measured by allowing them to recover in intracellular buffer before the addition of 100nM CaCl2 plus 2.5 µM CCCP (to block mitochondrial Ca^2+^ uptake). To mimic cytosolic ionic composition in these permeabilizing experiments, HEK293T cells were washed with high K^+^ intracellular buffer, containing 110 mM KCl, 10 mM NaCl, 0.5 mM K2HPO4, 5 mM succinate, 10 mM HEPES (pH 7.0 at 37 °C), supplemented with 5 mM EDTA or 1 mM EGTA. 100nM [Ca^2+^] was calculated using the Maxchelator program.

### Assessment of SERCA conformational changes

FRET imaging experiments were performed using an epifluorescence inverted microscope (DMIRE-2, Leica) with a PlanApo 40x oil immersion objective. The excitation light source was a high-speed scanning polychromator with a Xe lamp (C7773, Hamamatsu Photonics). An emission filter wheel was controlled by a Lambda-10 device (Sutter Instruments). Images were acquired with an ORCA-FLASH 4.0 camera (C11440-22CU) and Aquacosmos 2.6 software was used to control all devices. Both camera and software were from Hamamatsu Photonics. ImageJ software was used for analyses.

Cells were plated in 35 mm ibidi µ-dishes, and transfected with lipofectamine 2000. The SERCA2a conformation sensor was labelled with GFP in the N-terminus and tagRFP in the nucleotide domain. Transfections were performed using RG-SERCA and either empty vector, CFP-WT STIM1 or CFP-E152K-STIM1 (1 µg:0.5 µg DNA). Cells were perfused in a medium containing: 135 mM NaCl, 5 mM KCl, 1 mM MgCl2, 1.8 mM CaCl2, 10 mM Hepes, 10 mM glucose, pH 7.4. The Ca^2+-^free solution contained 1mM EGTA instead of 1.8 mM CaCl2. Cells were illuminated at 500 nm through an excitation filter (422/503/572 nm) and a triple-edge dichroic mirror (422/520/590 nm). FRET and donor emissions were alternately collected with emission filters 620/52 nm and 535/22 nm (center wavelength/bandwidth), respectively. A ratio of FRET image (donor excitation, acceptor emission) and donor image (GFP) (donor excitation, donor emission) was calculated, after subtracting background in both channels for its analysis. STIM1 expression was checked by excitation at 430 nm whilst emission was collected with emission filter 475/20 nm.

### Western blotting

Cells were washed in ice-cold PBS and lysed with RIPA buffer containing protease inhibitor cocktail at 4°C for 30 min. Cell lysates were centrifuged at 16,000g for 10 min at 4°C, and supernatants were collected. Cell lysates (100 µg) were loaded, separated in acrylamide SDS-PAGE gel (gradient 4-12%), and transferred to PVDF membrane. For experiments designed to evaluate trypsin secretion, the conditioned media were treated with trichloroacetic acid for protein precipitation and washed three times in acetone. Blots were incubated with anti-PRSS1 antibody (1:1000), anti-STIM1 antibody (1:1000) or anti-GAPDH antibody (1:2000) and probed with a peroxidase-conjugated anti-mouse antibody (1:2000) or anti-sheep secondary antibody (1:2000). An enhanced chemiluminescence system was used for visualization.

### Immunoprecipitation

Anti-GOK/STIM1 antibody was coupled to Protein G magnetic beads (Invitrogen) by incubating 2.5 µg antibody with the beads (30 µl) for 1 hour at 4°C. Cells were washed in ice-cold PBS and treated with lysis buffer containing protease inhibitor cocktail at 4°C for 30 min. Cell lysates were centrifuged at 14000 g for 30 min at 4°C after sonication, and supernatants were collected. Cell lysates (500 mg) were added into the beads coupled to the antibody and incubated under gently agitation overnight at 4°C. Precipitated proteins were eluted with SDS-sample buffer and incubated at 70°C for 10 min.

### STIM1 Oligomerization

1 µg of CFP-STIM1 constructs and 3 µg YFP-STIM1 constructs were co-transfected and cells were imaged 48 h after transfection. The oligomerization of STIM1 was followed by measuring the FRET signal between CFP- and YFP-STIM1. Cells were excited at 430 nm through a 455-nm dichroic mirror (455DRLP, Omega Optical), and emission was collected alternatively at 480 and 535 nm (480AF30 and 535DF25, Omega Optical).

### Bzipar assay

Cells were incubated with 100 µM of the trypsin substrate BZiPAR (Thermo Fisher Scientific, Waltham, MA) for 120 min. Cells were excited at 494 nm for 150 ms, and fluorescence emission was collected at 535 nm. A T510lpxrxt dichroic filter and an ET535/50 emission filter both from Chroma Technology (Bellows Falls, VT)] were used. Trypsin was added to confirm the increase in fluorescence signal when BZiPAR was present. The trypsin inhibitor benzamidine (5 ug/ml) was used to check specific trypsin response. The fluorescence (excitation 494-emission 535 nm) was measured from each cell to compare among conditions.

### Cytotoxicity assay

Cell toxicity was measured using the CellTox™ Green Cytotoxicity Assay kit (Promega), and results were normalized to living cell numbers as evaluated with the CellTiter 96^®^ AQueous One Solution Cell Proliferation Assay Kit (Promega) following the manufacturer’s instructions. Absorbance, which is proportional to living cell numbers, was measured at 490 nm. The cell cytotoxicity test is based on cellular membrane integrity using a cyanine dye with significant affinity for the DNA of dying cells. The dye properties are modified when fixed to DNA and fluorescence is enhanced. Emitted fluorescence (excitation: 512 nm, emission: 532 nm) is proportional to cell cytotoxicity and was measured for each experimental condition at different time points following the manufacturer’s instructions. Cytotoxicity results were normalized using cell titration data.

### Recombinant EF-SAM protein expression and purification

The T153I and E152K variants were separately introduced into pET-28a expression vectors using the Quikchange protocol (Agilent, Inc.). The pET-28a vectors were transformed into BL21 DE3 CodonPlus E. coli cells by heat shock, and liquid Luria-Bertani cultures were grown at 37°C until the optical density (600nm) reached 0.6. Protein expression was subsequently induced with 0.3 mM isopropyl beta-D-thiogalactopyranoside and allowed to progress overnight. The hexahistidine (His6)-tagged proteins of interest were captured from E. coli cell lysate using Ni-nitriloacetic acid agarose resin (Qiagen) under denaturing conditions according to the manufacturer’s guidelines. EF-SAM was refolded by dilution into 20mM Tris, 150mM NaCl, 5mM CaCl2, pH 8. Subsequently, the His6-tags were cleaved by thrombin digestion and a final size exclusion chromatography purification step using a Superdex 200 10/300 GL column linked with an AKTA FPLC (GE Healthcare, Inc.) was performed to achieve a purity of > 95%.

### Far-UV-CD

CD data were acquired on a Jasco J-815 CD spectrometer (Jasco, Inc.) collected in 1 nm increments (20 nm min-1) using a 0.1 cm path length cuvette, 8 s averaging time, and 1 nm bandwidth at 20 °C. Thermal melts were acquired by monitoring the change in the 225 nm CD signal as a function of temperature (i.e. 4-75 °C) using 0.1 cm cuvettes, 8 s averaging time, 1 nm bandwidth and 1 °C min-1 scan rate. Spectra were corrected for buffer contributions.

### SEC-MALS

MALS measurements were performed in-line with the size exclusion chromatography using a three angle (i.e. 45°, 90° and 135°) miniDawn light scattering instrument equipped with a 690nm laser and an Optilab rEX differential refractometer (Wyatt Technologies, Inc.). Molecular weight was calculated using ASTRA software (Wyatt Technologies, Inc.) based on Zimm plot analysis and using a protein refractive index increment, dndc −1 = 0.185 L g-1.

### Statistical analyses

Sets of functional analytic data were compared using analysis of variance (ANOVA) or a Student’s t-test. *, *P* < 0.05; ** *P* < 0.001; *** *P* < 0.0005; **** *P* <0.0001. Data shown were from at least four independent experiments.

Computational methodology is described in detail in Supp. materials and methods.

## ACKNOWLEDGEMENTS

We thank Dr. N. Demaurex (Geneva, Switzerland) for generously providing the D1ER plasmid, Drs. R. Y. Tsien (La Jolla, CA) and A. Palmer (La Jolla, CA) for providing the cameleon. We are grateful to M. Henry (Brest, France), A. Youinou (Brest, France) and L. Leroi (Brest, France) for technical support. We also thank C. Castelbou (Geneva, Switzerland) for technical assistance.

## Disclosure statement

The authors declare no conflict of interest.

## Funding information

M. Burgos was supported by the Association de Transfusion Sanguine et de Biogénétique Gaetan Saleun, the Spanish Ministry of Economy and Competitivity, reference BFU2015-69874-R and the Castilla-La Mancha Government, reference II-2018_11. R. Philippe and O. Mignen were supported by the French association Vaincre La Mucoviscidose and the Association de Transfusion Sanguine et de Biogénétique Gaetan Saleun. F. Antigny was supported by a post-doctoral grant from Aviesan (ITMO IHP). M. Frieden was supported by the Swiss National Foundation grant # 310030-141113. the Association des Pancréatites Chroniques Héréditaires and the Institut National de la Santé et de la Recherche Médicale (INSERM), France. This work was supported by CIHR, HSFC and NSERC operating grants to M.Ikura and an NSERC operating grant to P. Stathopulos.

## Author contributions

Project design and coordination were carried out by OM and MB. The writing group comprised OM and MB. Functional analyses were directed by OM. MB designed and performed time-lapse Ca^2+^, and BiPAR experiments with the assistance of RP. RP performed cytotoxicity experiments. FA and RP performed ER Ca^2+^ and FRET experiments under the supervision of MF. PD and PB performed SERCA co-immunoprecipitation experiments. CLM and FC were involved in the identification of patients and sample collection. Fibroblast isolation and culture were carried out by NL. JL, BD and MB designed and performed SERCA-STIM1 FRET experiments. PS and MI designed and performed Far-UV-CD and SECS-MALS experiments. WB, SM and WG performed computational experiments. TC, JMC and CF critically revised the manuscript and contributed to data interpretation. All authors approved the final manuscript and contributed critical revisions to its intellectual content.

## Abbreviations

EF-SAM: EF hand/Sterile Alpha Motif
STIM1: Stromal Interaction Molecule 1
ER: Endoplasmic Reticulum
SERCA: Sarco Endoplasmic Reticulum Ca2+ ATPase
SOCE: Store Operated Ca2+ Entry
CCK: Cholecystokinin
InsP3-R: Inositol Triphosphate Receptor
CC: Coiled-Coil
ICP: Idiopathic Chronic Pancreatitis
PMCA: Plasma Membrane Ca2+ ATPase
WT: wild-type
Ach: Acetylcholine
CCh: Carbachol
PLC: Phospholipase C
SOCIC: Store-Operated Ca2+ Influx Complex
FRET: Förster Resonance Energy Transfer
CD: Circular Dichroism
SEC-MALS: Size Exclusion Chromatography with in-line Multiangle Light Scattering
tBHQ: tert-Butylhydroquinone

## REFERENCES

Böhm, J., Chevessier, F., Maues De Paula, A., Koch, C., Attarian, S., Feger, C., Hantaï, D., Laforêt, P., Ghorab, K., Vallat, J.-M., et al. (2013). Constitutive activation of the calcium sensor STIM1 causes tubular-aggregate myopathy. Am. J. Hum. Genet. 92, 271–278.

Gerasimenko, J. V., Lur, G., Sherwood, M. W., Ebisui, E., Tepikin, A. V., Mikoshiba, K., Gerasimenko, O. V. and Petersen, O. H. (2009). Pancreatic protease activation by alcohol metabolite depends on Ca2+ release via acid store IP3 receptors. Proc. Natl. Acad. Sci. U. S. A. 106, 10758–10763.

Gerasimenko, J. V., Gryshchenko, O., Ferdek, P. E., Stapleton, E., Hébert, T. O. G., Bychkova, S., Peng, S., Begg, M., Gerasimenko, O. V. and Petersen, O. H. (2013). Ca2+ release-activated Ca2+ channel blockade as a potential tool in antipancreatitis therapy. Proc. Natl. Acad. Sci. U. S. A. 110, 13186–13191.

Gerasimenko, J. V., Gerasimenko, O. V. and Petersen, O. H. (2014). The role of Ca2+ in the pathophysiology of pancreatitis. J. Physiol. 592, 269–280.

Harris, E., Burki, U., Marini-Bettolo, C., Neri, M., Scotton, C., Hudson, J., Bertoli, M., Evangelista, T., Vroling, B., Polvikoski, T., et al. (2017). Complex phenotypes associated with STIM1 mutations in both coiled coil and EF-hand domains. Neuromuscul. Disord. NMD 27, 861–872.

Hedberg, C., Niceta, M., Fattori, F., Lindvall, B., Ciolfi, A., D’Amico, A., Tasca, G., Petrini, S., Tulinius, M., Tartaglia, M., et al. (2014). Childhood onset tubular aggregate myopathy associated with de novo STIM1 mutations. J. Neurol. 261, 870–876.

Hou, Z., Hu, Z., Blackwell, D. J., Miller, T. D., Thomas, D. D. and Robia, S. L. (2012). 2-Color calcium pump reveals closure of the cytoplasmic headpiece with calcium binding. PloS One 7, e40369.

Jousset, H., Frieden, M. and Demaurex, N. (2007). STIM1 knockdown reveals that store-operated Ca2+ channels located close to sarco/endoplasmic Ca2+ ATPases (SERCA) pumps silently refill the endoplasmic reticulum. J. Biol. Chem. 282, 11456–11464.

Kereszturi, E. and Sahin-Tóth, M. (2009). Intracellular autoactivation of human cationic trypsinogen mutants causes reduced trypsinogen secretion and acinar cell death. J. Biol. Chem. 284, 33392–33399.

Krüger, B., Albrecht, E. and Lerch, M. M. (2000). The role of intracellular calcium signaling in premature protease activation and the onset of pancreatitis. Am. J. Pathol. 157, 43–50.

Lacruz, R. S. and Feske, S. (2015). Diseases caused by mutations in ORAI1 and STIM1. Ann. N. Y. Acad. Sci. 1356, 45–79.

Liou, J., Kim, M. L., Heo, W. D., Jones, J. T., Myers, J. W., Ferrell, J. E. and Meyer, T. (2005). STIM is a Ca2+ sensor essential for Ca2+-store-depletion-triggered Ca2+ influx. Curr. Biol. CB 15, 1235–1241.

López, J. J., Jardín, I., Bobe, R., Pariente, J. A., Enouf, J., Salido, G. M. and Rosado, J. A. (2008). STIM1 regulates acidic Ca2+ store refilling by interaction with SERCA3 in human platelets. Biochem. Pharmacol. 75, 2157–2164.

Lur, G., Sherwood, M. W., Ebisui, E., Haynes, L., Feske, S., Sutton, R., Burgoyne, R. D., Mikoshiba, K., Petersen, O. H. and Tepikin, A. V. (2011). InsP₃receptors and Orai channels in pancreatic acinar cells: co-localization and its consequences. Biochem. J. 436, 231–239.

Manjarrés, I. M., Rodríguez-García, A., Alonso, M. T. and García-Sancho, J. (2010). The sarco/endoplasmic reticulum Ca(2+) ATPase (SERCA) is the third element in capacitative calcium entry. Cell Calcium 47, 412–418.

Maruyama, Y. and Petersen, O. H. (1994). Delay in granular fusion evoked by repetitive cytosolic Ca2+ spikes in mouse pancreatic acinar cells. Cell Calcium 16, 419–430.

Maruyama, Y., Inooka, G., Li, Y. X., Miyashita, Y. and Kasai, H. (1993). Agonist-induced localized Ca2+ spikes directly triggering exocytotic secretion in exocrine pancreas. EMBO J. 12, 3017–3022.

Masamune, A., Kotani, H., Sörgel, F. L., Chen, J.-M., Hamada, S., Sakaguchi, R., Masson, E., Nakano, E., Kakuta, Y., Niihori, T., et al. (2020). Variants That Affect Function of Calcium Channel TRPV6 Are Associated With Early-onset Chronic Pancreatitis: TRPV6 and pancreatitis. Gastroenterology.

Messenger, S. W., Falkowski, M. A. and Groblewski, G. E. (2014). Ca^2+^-regulated secretory granule exocytosis in pancreatic and parotid acinar cells. Cell Calcium 55, 369–375.

Misceo, D., Holmgren, A., Louch, W. E., Holme, P. A., Mizobuchi, M., Morales, R. J., De Paula, A. M., Stray-Pedersen, A., Lyle, R., Dalhus, B., et al. (2014). A dominant STIM1 mutation causes Stormorken syndrome. Hum. Mutat. 35, 556–564.

Morin, G., Bruechle, N. O., Singh, A. R., Knopp, C., Jedraszak, G., Elbracht, M., Brémond-Gignac, D., Hartmann, K., Sevestre, H., Deutz, P., et al. (2014). Gain-of-Function Mutation in STIM1 (P.R304W) Is Associated with Stormorken Syndrome. Hum. Mutat. 35, 1221–1232.

Nesin, V., Wiley, G., Kousi, M., Ong, E.-C., Lehmann, T., Nicholl, D. J., Suri, M., Shahrizaila, N., Katsanis, N., Gaffney, P. M., et al. (2014). Activating mutations in STIM1 and ORAI1 cause overlapping syndromes of tubular myopathy and congenital miosis. Proc. Natl. Acad. Sci. U. S. A. 111, 4197–4202.

Noury, J.-B., Böhm, J., Peche, G. A., Guyant-Marechal, L., Bedat-Millet, A.-L., Chiche, L., Carlier, R.-Y., Malfatti, E., Romero, N. B. and Stojkovic, T. (2017). Tubular aggregate myopathy with features of Stormorken disease due to a new STIM1 mutation. Neuromuscul. Disord. NMD 27, 78–82.

Nwokonko, R. M., Cai, X., Loktionova, N. A., Wang, Y., Zhou, Y. and Gill, D. L. (2017). The STIM-Orai Pathway: Conformational Coupling Between STIM and Orai in the Activation of Store-Operated Ca2+ Entry. Adv. Exp. Med. Biol. 993, 83–98.

Pallikkuth, S., Blackwell, D. J., Hu, Z., Hou, Z., Zieman, D. T., Svensson, B., Thomas, D. D. and Robia, S. L. (2013). Phosphorylated phospholamban stabilizes a compact conformation of the cardiac calcium-ATPase. Biophys. J. 105, 1812–1821.

Palmer, A. E., Jin, C., Reed, J. C. and Tsien, R. Y. (2004). Bcl-2-mediated alterations in endoplasmic reticulum Ca2+ analyzed with an improved genetically encoded fluorescent sensor. Proc. Natl. Acad. Sci. U. S. A. 101, 17404–17409.

Park, C. Y., Hoover, P. J., Mullins, F. M., Bachhawat, P., Covington, E. D., Raunser, S., Walz, T., Garcia, K. C., Dolmetsch, R. E. and Lewis, R. S. (2009). STIM1 clusters and activates CRAC channels via direct binding of a cytosolic domain to Orai1. Cell 136, 876–890.

Parry, D. A., Holmes, T. D., Gamper, N., El-Sayed, W., Hettiarachchi, N. T., Ahmed, M., Cook, G. P., Logan, C. V., Johnson, C. A., Joss, S., et al. (2016). A homozygous STIM1 mutation impairs store-operated calcium entry and natural killer cell effector function without clinical immunodeficiency. J. Allergy Clin. Immunol. 137, 955–957.e8.

Petersen, O. H. (2009). Ca2+ signaling in pancreatic acinar cells: physiology and pathophysiology. Braz. J. Med. Biol. Res. Rev. Bras. Pesqui. Medicas E Biol. 42, 9–16.

Petersen, O. H. (2015). Ca^2+^ signalling in the endoplasmic reticulum/secretory granule microdomain. Cell Calcium 58, 397–404.

Philippe, R., Antigny, F., Buscaglia, P., Norez, C., Becq, F., Frieden, M. and Mignen, O. (2015). SERCA and PMCA pumps contribute to the deregulation of Ca2+ homeostasis in human CF epithelial cells. Biochim. Biophys. Acta 1853, 892–903.

Picard, C., McCarl, C.-A., Papolos, A., Khalil, S., Lüthy, K., Hivroz, C., LeDeist, F., Rieux-Laucat, F., Rechavi, G., Rao, A., et al. (2009). STIM1 mutation associated with a syndrome of immunodeficiency and autoimmunity. N. Engl. J. Med. 360, 1971–1980.

Raraty, M., Ward, J., Erdemli, G., Vaillant, C., Neoptolemos, J. P., Sutton, R. and Petersen, O. H. (2000). Calcium-dependent enzyme activation and vacuole formation in the apical granular region of pancreatic acinar cells. Proc. Natl. Acad. Sci. U. S. A. 97, 13126–13131.

Ritchie, M. F., Samakai, E. and Soboloff, J. (2012). STIM1 is required for attenuation of PMCA-mediated Ca2+ clearance during T-cell activation. EMBO J. 31, 1123–1133.

Roos, J., DiGregorio, P. J., Yeromin, A. V., Ohlsen, K., Lioudyno, M., Zhang, S., Safrina, O., Kozak, J. A., Wagner, S. L., Cahalan, M. D., et al. (2005). STIM1, an essential and conserved component of store-operated Ca2+ channel function. J. Cell Biol. 169, 435–445.

Sampieri, A., Zepeda, A., Asanov, A. and Vaca, L. (2009). Visualizing the store-operated channel complex assembly in real time: identification of SERCA2 as a new member. Cell Calcium 45, 439–446.

Satoh, K., Matsu-Ura, T., Enomoto, M., Nakamura, H., Michikawa, T. and Mikoshiba, K. (2011). Highly cooperative dependence of sarco/endoplasmic reticulum calcium ATPase SERCA2a pump activity on cytosolic calcium in living cells. J. Biol. Chem. 286, 20591–20599.

Shen, W.-W., Frieden, M. and Demaurex, N. (2011). Local cytosolic Ca2+ elevations are required for stromal interaction molecule 1 (STIM1) de-oligomerization and termination of store-operated Ca2+ entry. J. Biol. Chem. 286, 36448–36459.

Sofia, V. M., Da Sacco, L., Surace, C., Tomaiuolo, A. C., Genovese, S., Grotta, S., Gnazzo, M., Ciocca, L., Petrocchi, S., Alghisi, F., et al. (2016). Extensive molecular analysis suggested the strong genetic heterogeneity of idiopathic chronic pancreatitis. Mol. Med. Camb. Mass 22, 300–309.

Son, A., Ahuja, M., Schwartz, D. M., Varga, A., Swaim, W., Kang, N., Maleth, J., Shin, D. M. and Muallem, S. (2019). Ca2+ Influx Channel Inhibitor SARAF Protects Mice From Acute Pancreatitis. Gastroenterology.

Stathopulos, P. B., Li, G.-Y., Plevin, M. J., Ames, J. B. and Ikura, M. (2006). Stored Ca2+ depletion-induced oligomerization of stromal interaction molecule 1 (STIM1) via the EF-SAM region: An initiation mechanism for capacitive Ca2+ entry. J. Biol. Chem. 281, 35855–35862.

Stathopulos, P. B., Zheng, L., Li, G.-Y., Plevin, M. J. and Ikura, M. (2008). Structural and mechanistic insights into STIM1-mediated initiation of store-operated calcium entry. Cell 135, 110–122.

Szabó, A., Radisky, E. S. and Sahin-Tóth, M. (2014). Zymogen activation confers thermodynamic stability on a key peptide bond and protects human cationic trypsin from degradation. J. Biol. Chem. 289, 4753–4761.

Szmola, R. and Sahin-Tóth, M. (2007). Chymotrypsin C (caldecrin) promotes degradation of human cationic trypsin: identity with Rinderknecht’s enzyme Y. Proc. Natl. Acad. Sci. U. S. A. 104, 11227–11232.

Vaca, L. (2010). SOCIC: the store-operated calcium influx complex. Cell Calcium 47, 199–209.

Vaeth, M., Maus, M., Klein-Hessling, S., Freinkman, E., Yang, J., Eckstein, M., Cameron, S., Turvey, S. E., Serfling, E., Berberich-Siebelt, F., et al. (2017). Store-Operated Ca2+ Entry Controls Clonal Expansion of T Cells through Metabolic Reprogramming. Immunity 47, 664–679.e6.

Waldron, R. T., Chen, Y., Pham, H., Go, A., Su, H.-Y., Hu, C., Wen, L., Husain, S. Z., Sugar, C. A., Roos, J., et al. (2019). The Orai Ca2+ channel inhibitor CM4620 targets both parenchymal and immune cells to reduce inflammation in experimental acute pancreatitis. J. Physiol. 597, 3085–3105.

Wen, L., Voronina, S., Javed, M. A., Awais, M., Szatmary, P., Latawiec, D., Chvanov, M., Collier, D., Huang, W., Barrett, J., et al. (2015). Inhibitors of ORAI1 Prevent Cytosolic Calcium-Associated Injury of Human Pancreatic Acinar Cells and Acute Pancreatitis in 3 Mouse Models. Gastroenterology 149, 481–492.e7.

Wu, M. M., Buchanan, J., Luik, R. M. and Lewis, R. S. (2006). Ca2+ store depletion causes STIM1 to accumulate in ER regions closely associated with the plasma membrane. J. Cell Biol. 174, 803–813.

Yuan, J. P., Kim, M. S., Zeng, W., Shin, D. M., Huang, G., Worley, P. F. and Muallem, S. (2009). TRPC channels as STIM1-regulated SOCs. Channels Austin Tex 3, 221–225.

Zhu, Z.-D., Yu, T., Liu, H.-J., Jin, J. and He, J. (2018). SOCE induced calcium overload regulates autophagy in acute pancreatitis via calcineurin activation. Cell Death Dis. 9, 50.

Zou, W.-B., Tang, X.-Y., Zhou, D.-Z., Qian, Y.-Y., Hu, L.-H., Yu, F.-F., Yu, D., Wu, H., Deng, S.-J., Lin, J.-H., et al. (2018). SPINK1, PRSS1, CTRC, and CFTR Genotypes Influence Disease Onset and Clinical Outcomes in Chronic Pancreatitis. Clin. Transl. Gastroenterol. 9, 204.

